# Carboxyl Terminal Domain Missense Mutations Alter Distinct Gating Properties of the Cardiac Sodium Channel

**DOI:** 10.1101/2025.05.09.653108

**Authors:** Akshay Sharma, Christopher Marra, Nomon Mohammad, Vasilisa Iatckova, Lillian Lawrence, Mitchell Goldfarb

## Abstract

Voltage-gated sodium channels undergo reversible voltage/time-dependent transitions from closed to open and inactivated states. The voltage setpoints and efficiency of cardiac sodium channel Na_v_1.5 state transitions are crucial for tuning the initiation and conduction of myocardial action potentials. The channel’s cytoplasmic carboxyl-terminal domain (CTD) regulates gating by intramolecular interactions and by serving as a hub for the binding of accessory proteins. We have investigated the roles of the CTD in intrinsic and FGF homologous factor (FHF)-modulated Na_v_1.5 gating through structure-guided CTD subdomain mutagenesis. The EF-hand module within the CTD was found to exert the most profound effects on channel gating, strongly influencing voltage-dependence of inactivation and activation, accelerating inactivation from the closed state, decelerating inactivation from the open state, minimizing persistent sodium current, and serving as the binding domain for FHF proteins. Na_v_1.5^D1788K^ bearing a missense mutation in the EF-hand motif displayed a depolarizing shift in voltage dependence of activation and generated greatly enhanced persistent sodium current without altering the voltage dependence of channel inactivation. Reciprocally, Na_v_1.5^L1861A^ bearing a different missense mutation in the EF-hand underwent closed-state inactivation at more negative membrane potential and at an accelerated rate, but did not display other phenotypes associated with CTD deletion. Na_v_1.5^V1776A/T1778A^ bearing mutations in the juxtamembrane region between the EF-hand and the channel pore helices displayed wild-type intrinsic gating properties, while FHF modulation of inactivation gating was impaired. Our channel physiology studies together with prior structural data suggest that the voltage and rate of channel inactivation from the closed state are governed by an intramolecular hydrophobic interaction of the CTD EF-hand with the cytoplasmic inactivation loop helix and the extension of this binding interface upon FHF-induced restructuring of the juxtamembrane region, while a distinct CTD intramolecular electrostatic interaction modulates voltage-dependent activation and minimizes persistent sodium current.

## Introduction

Action potentials in excitable cells are driven by the precise conformational tuning of voltage-gated sodium channels (VGSCs). Mutations in genes encoding the conducting alpha subunits of VGSCs or channel-binding accessory subunits cause a wide range of clinical dysfunctions, including early-onset epilepsies (O’Malley and Isom, 2015; Siekierska et al., 2016; Catterall, 2018; Fry et al., 2021; Talwar and Hammer, 2021), cardiac arrhythmias (Kapplinger et al., 2010; Wilde and Amin, 2018), neuromuscular disorders (Cannon, 2018), spinocerebellar ataxia (Groth and Berman, 2018), and pain disorders (Dib-Hajj and Waxman, 2019). VGSC alpha subunits have four subdomains constituting the pore and voltage sensors (Figure 1A) that actuate intrinsic mechanisms for voltage- and time-dependent transitions among their three principle states: closed-state (otherwise termed available), open-state (otherwise termed activated), and inactivated-state (Catterall, 2014). Channel activation results from depolarization-dependent outward movement of positively charged S4 helices on each of the subdomains (Bosmans et al., 2008), while channel inactivation results from depolarization-dependent movement of the S4 helix on subdomain IV along with the docking of the cytoplasmic loop between subdomains III and IV onto the internal face of the channel pore (West et al., 1992; Sheets et al., 1999; Capes et al., 2013; Burel et al., 2017). The stability of inactivation loop docking minimizes the probability of single-channel reopening and, hence, limits so-called persistent or late current in the depolarized cell (Sakmann et al., 2000).

**Figure 1:**
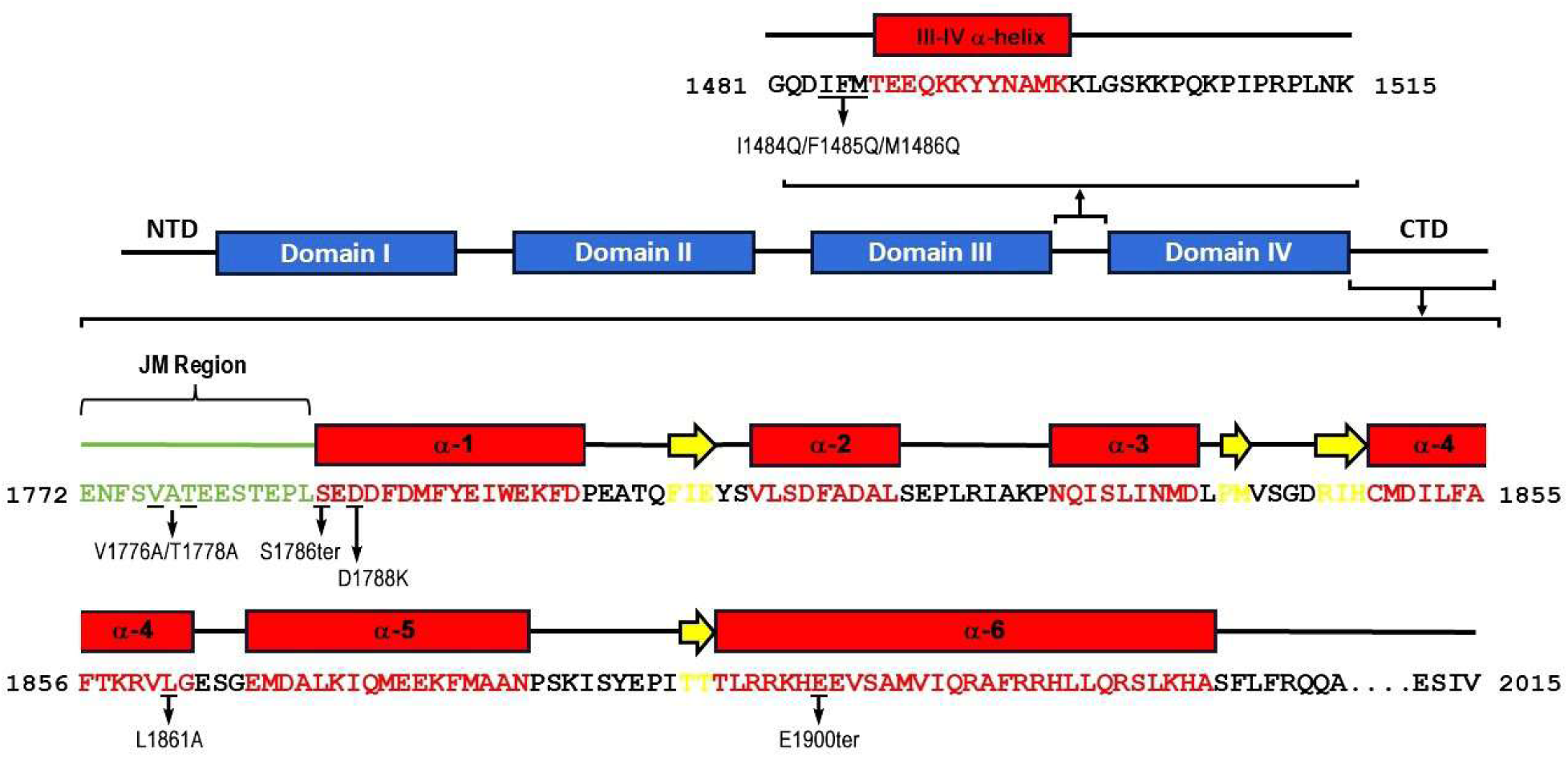
Na_v_1.5 domain map and amino acid sequences in the CTD and the III-IV inactivation loop. The four pseudotetrameric domains (Domain I through Domain IV) are colored blue. The alpha helices within the CTD and the III-IV inactivation loop are colored red, residues in beta strands are colored yellow, and the JM region within the CTD is colored green. Amino acid residues targeted for missense or nonsense mutations in this study are indicated.

The cytoplasmic C-terminal domains (CTD) of a VGSC alpha subunit bears a short juxtamembrane (JM) segment, an EF-hand fold consisting of five alpha helices, a long α6 helix, and a tail segment (Fig 1A) (Wang et al., 2012). The CTDs of the nine mammalian VGSC alpha subunits (Na_v_1.1 – Na_v_1.9) display extensive sequence conservation throughout the JM (85% identity), EF-hand (76% identity) and α6 helix (73% identity) segments, while the tail segments are highly divergent in sequence and length (Catterall, 2014). A role for the CTD in determining the voltage dependence of channel inactivation was first revealed by examining the gating properties of reciprocal Na_v_1.5(Na_v_1.2-CTD) and Na_v_1.2(Na_v_1.5-CTD) chimeric channels (Mantegazza et al., 2001). More recently identified arrhythmia-associated mutations altering the CTD of cardiac Na_v_1.5 have revealed roles for the EF-hand and α6 CTD segments in setting the voltage range for channel inactivation and in minimizing the late sodium current (Tan et al., 2007; Ziyadeh-Isleem et al., 2014; Musa et al., 2015; Yan et al., 2017a). The physical basis for CTD functionality is still uncertain, as the CTD is unresolved in the structures of full-length inactivated VGSCs (Yan et al., 2017b; Pan et al., 2018; Fan et al., 2023). However, a VGSC structurally resolved in the closed state showed an interaction between the EF-hand and the III-IV inactivation loop helix, suggesting that such an interaction may delay the transition to the inactivated state upon membrane depolarization (Shen et al., 2017). Indeed, various mutations in the EF-hand cause a hyperpolarizing shift in the voltage dependence of Na_v_1.5 inactivation (Cormier et al., 2002; Tan et al., 2007; Ziyadeh-Isleem et al., 2014; Gardill et al., 2018; Gade et al., 2020).

Fibroblast growth factor homologous factors (FHFs) are small cytoplasmic proteins that bind to mammalian VGSC CTDs and modulate channel inactivation (Goldfarb, 2024), thereby playing essential roles in central (Goldfarb et al., 2007; Laezza et al., 2007; Shakkottai et al., 2009; Venkatesan et al., 2014) and peripheral (Barbosa et al., 2017; Yang et al., 2017; Marra et al., 2023) neuron excitability as well as cardiomyocyte excitation and conduction (Wang et al., 2011a; Park et al., 2016; Wang et al., 2017; Chakouri et al., 2022). While binding of FHFs to the EF-hand domain of VGSC CTDs is essential for FHF function (Goetz et al., 2009), the mechanism by which VGSC:FHF interaction modulates steady-state inactivation is completely unknown, in large measure due to our still incomplete understanding of VGSC-intrinsic gating.

Here, we have employed structure-guided truncations and missense mutations in the CTD of Na_v_1.5 to further delineate the roles of CTD subdomains in intrinsic and FHF-modulated channel gating. Different missense mutations in the EF-hand segment of the CTD exerted distinct effects on persistent current and the gating of activation and inactivation. Furthermore, missense mutations in the JM segment had no observable effect on channel-intrinsic current density, channel gating, or FHF binding, while significantly impairing FHF modulation of inactivation gating, thereby defining the JM region as an FHF effector domain.

## Materials and methods

### Plasmids for mammalian expression

Human Na_v_1.5 cDNA used in this study corresponds to transcript variant 2 (accession NM 000335.5) that lacks codon Gln-1070 in the DII-DIII cytoplasmic loop due to alternative splicing. Hence, for all figures, tables, and text, the numbering of residues for the carboxyl-terminal half of Na_v_1.5 is one less than is more typically reported in the literature. This Na_v_1.5 cDNA in expression vector pCDNA3.1 and tetrodotoxin-resistant mouse Na_v_1.6^Y371S^ in vector pIRESneo3 (Dover et al., 2010) were mutated by Quickchange DNA synthesis using Pfu Turbo DNA polymerase (Agilent), and desired mutations in the absence of off-target mutations were confirmed by DNA sequencing (Eton Biosciences). FHF2B and FHF1B^R52H^ cDNAs cloned into bicistronic pIRES-ZsGreen1 vector (Clontech) were described previously (Dover et al., 2010; Siekierska et al., 2016).

### VGSC and FHF expression in Neuro2A cells

Channel-bearing plasmid and pIRES-ZsGreen (with or without FHF cDNA insertion) were co-transfected at a 2:1 ratio in Neuro2A cells using Lipofectamine 2000 (Invitrogen). Transfected cells were then trypsinized, plated onto poly-D-lysine treated glass coverslips, and used for electrophysiological recordings after 18-24 hours. When cells were to be used for protein extraction, transfer to coverslips was omitted.

### Voltage Clamp Sodium Current Recordings

All voltage clamp recordings were conducted using a MultiClamp700 Amplifier, Digidata 1440 analog/digital converter, and Clampex10 software (Molecular Devices). Cells were placed in the recording chamber, viewed with a Nikon Eclipse microscope and temperature was maintained at 25°C by an in-line heater (Warner Instruments). Cell-bearing coverslips were transferred to recording chamber containing carbogen-buffered bath solution (109mM NaCl, 26mM NaHCO3, 4.7mM KCl, 11mM glucose, 1.2mM MgCl2, 2mM CaCl2, 0.2mM CdCl2, 3mM myoinositol, 2mM Na pyruvate, 10mM HEPES pH 7.2) at 25 °C. For cells that would typically generate sodium currents in excess of 2 nA in the above bath solution, low-sodium bath solution (29mM NaCl, 80mM choline Cl, 26mM NaHCO3, 4.7mM KCl, 11mM glucose, 1.2mM MgCl2, 2mM CaCl2, 0.2mM CdCl2, 3mM myoinositol, 2mM Na pyruvate, 10mM HEPES pH 7.2) was employed. All extracellular bath solutions were supplemented with 0.3 μM tetrodotoxin to inhibit low-level Neuro2A endogenous sodium channels. Green fluorescent cells were whole-cell patched with pipettes filled with intracellular pipette solution containing 104mM CsCl, 50mM tetraethylamine chloride, 10mM HEPES pH 7.2, 5mM glucose, 2mM MgCl2, 10mM EGTA, 2mM Na2ATP, 0.2mM NaGTP and having 1–2 MΩ resistance. For all recording protocols, the voltage-gated current was isolated during data acquisition by P/N leak subtraction (N = 6). Current data were sampled at 10 or 20 kHz. All recording data were analyzed using Clampfit software.

### Voltage-clamp recording protocols and data analysis

Voltage dependence of activation and steady-state inactivation for Na_v_1.5 derivatives: A 31-sweep protocol consisted of −150 mV holding potential, a first 100 msec step at −150 + 5n mV (n = 0 to 30) followed by a 10 msec −10 mV step. Peak sodium currents from the first voltage step of all sweeps were used to calculate voltage dependence of channel activation, peak currents from the subsequent 10 mV step of all sweeps were used to calculate voltage dependence of steady-state inactivation. Peak conductances at each voltage were used to determine V1/2 and slope factor k by fitting to the Boltzmann equation: g(V) = g_max_/(1 + e^(V1/2 – V)/k^).

Voltage dependence for activation of inactivation-defective Na_v_1.5 mutant variants was assayed using a 19-sweep protocol consisting of −150 mV holding potential and step depolarization to −80 + 5n mV (n = 0 to 18), and the mean sodium current 25-30 msec after each depolarization was used to calculate voltage dependence of channel activation using the Boltzmann equation shown above.

Voltage dependence of activation and steady-state inactivation for Na_v_1.6^TTXr^ derivatives: A 26-sweep protocol consisted of −120 mV holding potential, a first 100 msec step at −120 + 5n mV (n = 0 to 25) followed by a 10 msec 0 mV step. Peak sodium currents from the first voltage step of all sweeps were used to calculate voltage dependence of channel activation, peak currents from the subsequent 0 mV step of all sweeps were used to calculate voltage dependence of steady-state inactivation using the Boltzmann equation shown above.

Rate of closed-state inactivation at −70 mV for Na_v_1.5 derivatives: An 11-sweep protocol consisted of − 150 mV holding potential, a first step at −70 mV for t msec (t = 0 to 10) followed by a 10 msec 0 mV step. Peak sodium currents from the 0 mV step for each sweep were used to calculate the rate constant tau of closed-state inactivation using the exponential decay equation: I(t) = I(0) * e^-t/tau^.

Rate of open-state inactivation: The decay phase of sodium current generated by step depolarization from −150 mV to −20 mV was used to calculate the rate constant tau of open-state inactivation using the exponential decay equation above.

Magnitude of persistent sodium current for Na_v_1.5 derivatives: A protocol of five replica sweeps consisted of −150 mV holding potential and depolarization to −10 mV for 50 msec. The replica sweeps were averaged, and persistent current measured after 30 msec of depolarization.

All calculated values from electrophysiological data were expressed as mean +/- standard error. Statistical significance between experimental and control groups was calculated using two-tailed unpaired Student’s T-test.

### Co-immunoprecipitation and Immunoblotting

Cells were lysed in Triton lysis buffer (20mM Tris pH 7.4, 137mM NaCl, 2mM EDTA, 25 mM β-glycerophosphate, 2mM Na pyrophosphate, 1 mM Na orthovanadate, 10% glycerol, 1% Triton X-100, 1 mM phenylmethanesulfanyl fluoride, 10 μg/ml aprotinin, 10 μg/ml leupeptin) and clarified by microfuge centrifugation at 15,000 rpm for ten minutes. 300 μg clarified lysate was adjusted to 0.5 ml in Triton lysis buffer and immunoprecipitated overnight at 4°C with 1.5 μg of rabbit anti-FHF2 (Schoorlemmer and Goldfarb, 2002), and immunoprecipitates were then captured by mixing with Protein G Sepharose. Immunoprecipitates were eluted from beads with Laemmli sample buffer (125 mM Tris pH 6.9, 2% sodium dodecyl sulphate [SDS], 2.5 M β-mercaptoethanol, 5 mg/ml bromophenol blue)at 60°C and electrophoresed through a 4-20% gradient polyacrylamide gel in presence of 0.1% SDS and 100 mM β-mercaptoethanol. Aliquots of clarified lysates were mixed with 3 volumes 2x Laemlli sample buffer and electrophoresed separately under same conditions. Proteins in gels were electrotransfered to polyvinylidene fluoride membrane, blocked with 10% horse serum, and probed with either anti-panNav mouse monoclonal antibody (Millipore Sigma) or rabbit anti-FHF2 antibodies, then incubated with secondary peroxidase-conjugated antibodies, followed by enhanced chemiluminescence detection.

### Statistical analysis

All calculated values for electrophysiological data and binding assays are expressed as mean +/- standard error. Statistical significance P values for comparison of two experimental groups were calculated using two-tailed unpaired Student’s t-test.

## Results

### Na_v_1.5^S1786ter^ Lacking the CTD EF-hand Motif Displays Altered Peak and Persistent Sodium Current and Setpoints for Channel Inactivation and Activation

Nonsense and missense mutations have been employed previously to explore the functionality of segments within the CTD of VGSCs, with the choice of mutations often focused on disease-associated variants. While this approach is of clinical value, such mutations may alter channel function by generating misfolded protein segments. In an attempt to obviate this shortcoming, we generated VGSC truncations guided by the known structural domains within the Na_v_1.5 CTD (Wang et al., 2012) (Figure 1). The nonsense mutation Na_v_1.5^E1900ter^ deletes the channel tail and the α6 helix, which harbors the IQ motif responsible for calmodulin binding. The nonsense mutation Na_v_1.5^S1786ter^ further deletes the helical EF-hand fold, while sparing the short juxtamembrane (JM) segment. Wild-type and truncated channels were expressed in Neuro2A cells, and voltage-clamp protocols were used to assess a cohort of channel properties: peak sodium current density, voltage-dependence of steady-state inactivation, closed- and open-state inactivation rates, voltage dependence of activation, and persistent current.

All properties of Na_v_1.5^E1900ter^ were statistically indistinguishable from those of wild-type Na_v_1.5 except for a small but significant increase in persistent current (Figure 2a,d,e; Table 1). The effect of this deletion on persistent current likely reflects the loss of the calmodulin-binding motif within the α6 helix (Yan et al., 2017a). By contrast, Na_v_1.5^S1786ter^ had greatly altered properties, including ∼10-fold reduction in peak current density (Table 1), a 13.7 mV hyperpolarizing shift in voltage-dependent inactivation (Figure 2a; Table 1), a 4-fold increase in the rate of closed-state inactivation at −70mV (Figure 2b; Table 1), a 2-fold decrease in the rate of open-state inactivation at −20 mV (Figure 2c; Table 1), and a 13-fold increase in the persistent:peak sodium current ratio (Figure 2d, Table 1).

**Figure 2.**
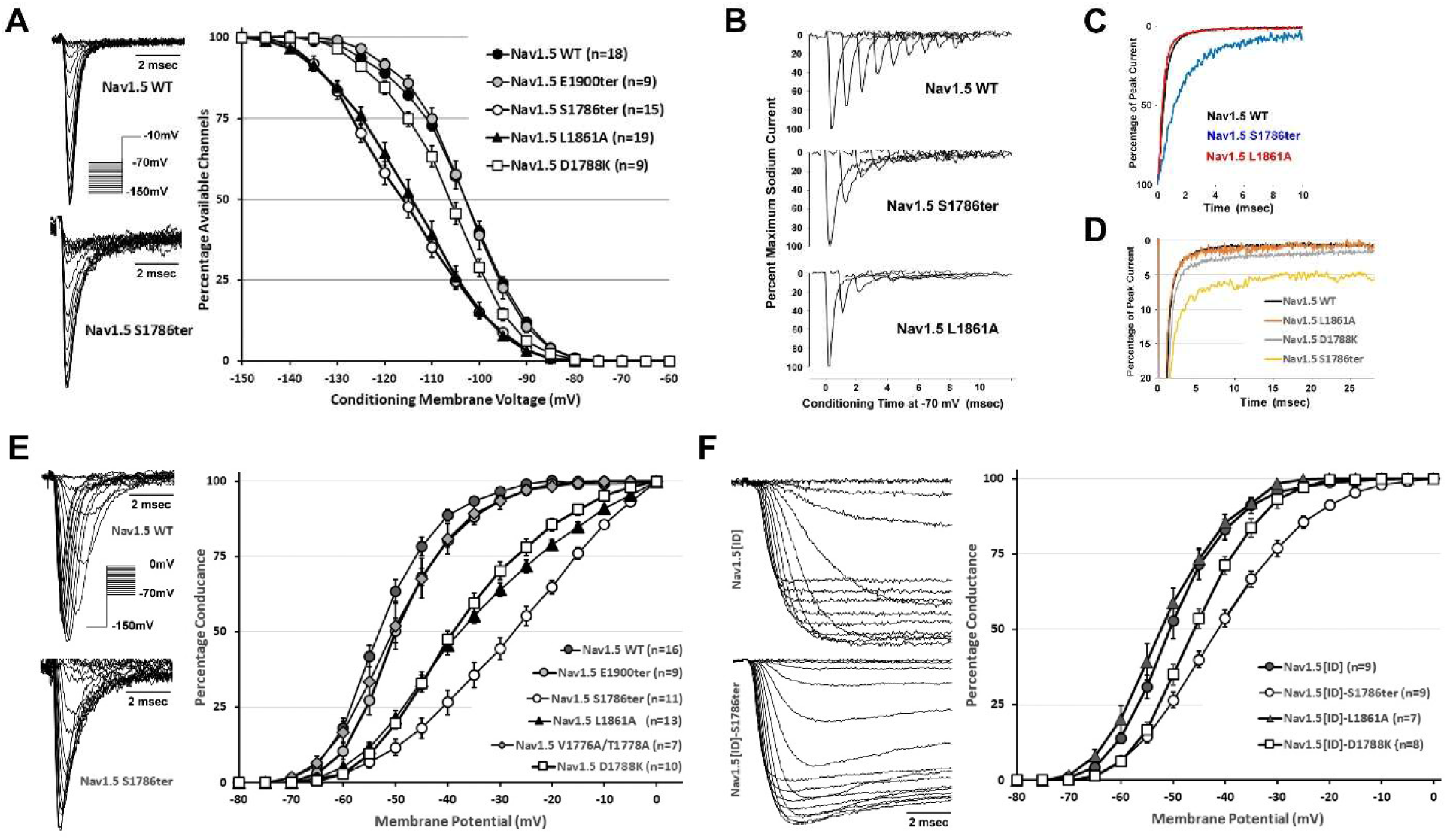
Mutations targeting the Na_v_1.5 CTD lead to diverse alterations of channel properties. **(A)** Left: Exemplar voltage-dependent sodium current traces for Na_v_1.5^wildtype^ and Na_v_1.5^S1786ter^ upon depolarization to −10 mV following 100 msec conditioning at test voltages. Right: Voltage-dependent steady-state inactivation curves show hyperpolarizing shifts for Na_v_1.5^S1786ter^ and Na_v_1.5^L1861A^. **(B)** Na_v_1.5^S1786ter^ and Na_v_1.5^L1861A^ undergo faster closed-state inactivation at −70mV compared to Na_v_1.5^wildtype^. Exemplar ensemble sodium current traces for depolarization to −10 mV following conditioning at −70 mV for 0-10 msec. **(C)** Exemplar traces displaying the decreased rate of Na_v_1.5^S1786ter^ open-state inactivation at −20 mV. The rate of Na_v_1.5^L1861A^ open-state inactivation is indistinguishable from Na_v_1.5^wildtype^. **(D)** Exemplar traces showing persistent currents at −10 mV for Na_v_1.5^wildtype^ and Na_v_1.5 mutant channels. **(E)** Left: Exemplar voltage-dependent activation traces for Na_v_1.5^wildtype^ and Na_v_1.5^S1786ter^. Right: Voltage-dependent activation curves. Na_v_1.5^S1786ter^, Na_v_1.5^D1788K^ and Na_v_1.5^L1861A^ display depolarizing shifts in the voltage-dependence of activation, while Na_v_1.5^E1900ter^ and Na_v_1.5^V1776A/T1778A^ do not. **(F)** Left: Exemplar voltage-dependent activation traces for Na_v_1.5[ID] and Na_v_1.5[ID]^S1786ter^. Right: Voltage-dependent activation curves showing the voltage-dependence of activation for Na_v_1.5[ID] mutants. Na_v_1.5[ID]^S1786ter^ and Na_v_1.5[ID]^D1788K^ showed a depolarized voltage- dependence of activation compared to Na_v_1.5[ID], while Na_v_1.5[ID]^L1861A^ does not. For all assays, the number of cells (n) measured for each transfection is indicated within the figure panels.

**Table 1.**
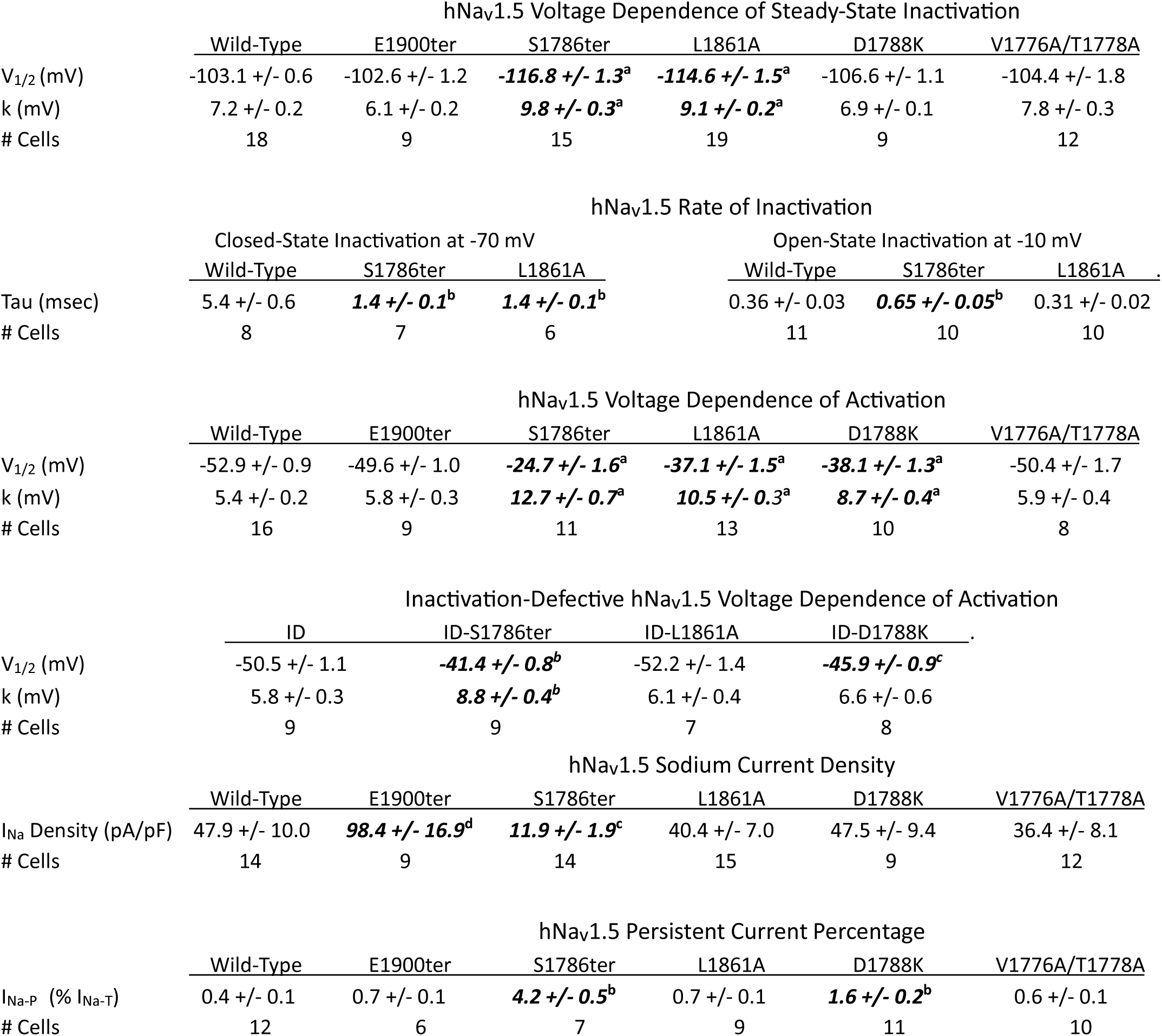
Current and gating properties of hNa_v_1.5 and mutant derivatives. Parameter values for mutants that differ significantly by Student t-Test from those of wild-type hNa_v_1.5 are shown in bold italics. ^a^P<0.000001, ^b^P<0.0001, ^c^P<0.004, ^d^P<0.025

| hNav1.5 Voltage Dependence of Steady-State Inactivation |  |  |  |  |  |  |
| --- | --- | --- | --- | --- | --- | --- |
|  | Wild-Type | E1900ter | S1786ter | L1861A | D1788K | V1776A/T1778A |
| V<sub>1/2</sub> (mV) | -103.1 +/- 0.6 | -102.6 +/- 1.2 | **-116.8 +/- 1.3<sup>a</sup>** | **-114.6 +/- 1.5<sup>a</sup>** | -106.6 +/- 1.1 | -104.4 +/- 1.8 |
| k (mV) | 7.2 +/- 0.2 | 6.1 +/- 0.2 | **9.8 +/- 0.3<sup>a</sup>** | **9.1 +/- 0.2<sup>a</sup>** | 6.9 +/- 0.1 | 7.8 +/- 0.3 |
| # Cells | 18 | 9 | 15 | 19 | 9 | 12 |

| hNav1.5 Rate of Inactivation |  |  |  |  |  |  |
| --- | --- | --- | --- | --- | --- | --- |
|  | Closed-State Inactivation at -70 mV |  |  | Open-State Inactivation at -10 mV |  |  |
|  | Wild-Type | S1786ter | L1861A | Wild-Type | S1786ter | L1861A |
| Tau (msec) | 5.4 +/- 0.6 | **1.4 +/- 0.1<sup>b</sup>** | **1.4 +/- 0.1<sup>b</sup>** | 0.36 +/- 0.03 | **0.65 +/- 0.05<sup>b</sup>** | 0.31 +/- 0.02 |
| # Cells | 8 | 7 | 6 | 11 | 10 | 10 |

| hNav1.5 Voltage Dependence of Activation |  |  |  |  |  |  |
| --- | --- | --- | --- | --- | --- | --- |
|  | Wild-Type | E1900ter | S1786ter | L1861A | D1788K | V1776A/T1778A |
| V<sub>1/2</sub> (mV) | -52.9 +/- 0.9 | -49.6 +/- 1.0 | **-24.7 +/- 1.6<sup>a</sup>** | **-37.1 +/- 1.5<sup>a</sup>** | **-38.1 +/- 1.3<sup>a</sup>** | -50.4 +/- 1.7 |
| k (mV) | 5.4 +/- 0.2 | 5.8 +/- 0.3 | **12.7 +/- 0.7<sup>a</sup>** | **10.5 +/- 0.3<sup>a</sup>** | **8.7 +/- 0.4<sup>a</sup>** | 5.9 +/- 0.4 |
| # Cells | 16 | 9 | 11 | 13 | 10 | 8 |

| Inactivation-Defective hNav1.5 Voltage Dependence of Activation |  |  |  |  |
| --- | --- | --- | --- | --- |
|  | ID | ID-S1786ter | ID-L1861A | ID-D1788K |
| V<sub>1/2</sub> (mV) | -50.5 +/- 1.1 | **-41.4 +/- 0.8<sup>b</sup>** | -52.2 +/- 1.4 | **-45.9 +/- 0.9<sup>c</sup>** |
| k (mV) | 5.8 +/- 0.3 | **8.8 +/- 0.4<sup>b</sup>** | 6.1 +/- 0.4 | 6.6 +/- 0.6 |
| # Cells | 9 | 9 | 7 | 8 |

| hNav1.5 Sodium Current Density |  |  |  |  |  |  |
| --- | --- | --- | --- | --- | --- | --- |
|  | Wild-Type | E1900ter | S1786ter | L1861A | D1788K | V1776A/T1778A |
| I<sub>Na</sub> Density (pA/pF) | 47.9 +/- 10.0 | **98.4 +/- 16.9<sup>d</sup>** | **11.9 +/- 1.9<sup>c</sup>** | 40.4 +/- 7.0 | 47.5 +/- 9.4 | 36.4 +/- 8.1 |
| # Cells | 14 | 9 | 14 | 15 | 9 | 12 |

TABLE 1. Current and Gating Properties of hNav1.5 and mutant derivatives
| hNav1.5 Persistent Current Percentage |  |  |  |  |  |  |
| --- | --- | --- | --- | --- | --- | --- |
|  | Wild-Type | E1900ter | S1786ter | L1861A | D1788K | V1776A/T1778A |
| I<sub>Na-P</sub> (% I<sub>Na-T</sub>) | 0.4 +/- 0.1 | 0.7 +/- 0.1 | **4.2 +/- 0.5<sup>b</sup>** | 0.7 +/- 0.1 | **1.6 +/- 0.2<sup>b</sup>** | 0.6 +/- 0.1 |
| # Cells | 12 | 6 | 7 | 9 | 11 | 10 |

Na_v_1.5^S1786ter^ also displayed an apparent 28 mV depolarizing shift in the voltage dependence of channel activation (Na_v_1.5^wildtype^ V_1/2_ = −52.9 mV vs. Na_v_1.5^S1786ter^ V_1/2_ = −24.7 mV) (Figure 2e; Table 1). This voltage shift could reflect an effect of CTD truncation on voltage sensor movements required to open the channel. Alternatively, the far faster inactivation of mutant channels from the closed state may reduce peak current amplitude at less depolarized voltages, thereby right-shifting the measured voltage at which most or all channels can open. To distinguish between these mechanisms, we engineered inactivation-defective (ID) variants of full-length and CTD-truncated channels by mutating the IlePheMet motif in the III-IV inactivation loop (Figure 1) (West et al., 1992). Indeed, preventing inactivation largely returned voltage dependence of activation for the CTD-truncated channel towards wild-type levels, demonstrating an effect of accelerated inactivation on the apparent voltage dependence of activation. However, activation of Na_v_1.5[ID]^S1786ter^ still displayed a statistically significant depolarizing shift (Na_v_1.5[ID]^S1786ter^ V_1/2_ = −41.4 +/- 0.8 mV) compared to full-length Na_v_1.5[ID] (V_1/2_ = −50.5 +/- 1.2 mV) (Figure 2f; Table 1), demonstrating that the CTD does indeed modulate the intrinsic activation mechanism.

Other laboratories have analyzed C-terminal truncations of Na_v_1.5. Na_v_1.5^S1884ter^ deletes the long α6 helix along with a structured linker between α6 and the EF-hand domain. Na_v_1.5^S1884ter^ exhibited an 11 mV hyperpolarizing shift in voltage dependence of inactivation, a substantial reduction in peak current density, and more elevated persistent:peak sodium current ratio (Cormier et al., 2002). It is not clear whether these effects, which trend towards the phenotype of Na_v_1.5^S1786ter^, reflect a functionality of the linker region between EF-hand and α6 segments or a destabilization of the EF-hand by deletion of the C-terminally adjacent residues. Separate studies have analyzed arrhythmia-associated frameshift (fs) mutations of Na_v_1.5 at Arg-1859 or Leu-1820, which truncate the CTD within the EF-hand segment (Tan et al., 2007; Ziyadeh-Isleem et al., 2014). The phenotype of these truncated channels includes changes in voltage-dependence and rate of channel inactivation, reduced current density and enhanced persistent current comparable to the degrees of Na_v_1.5^S1786ter^ reported here.

### EF-hand Missense Mutant Na_v_1.5^L1861A^ Phenocopies the Closed-State Inactivation Gating Abnormality of _Na_v_1.5_S1786ter

The structure of the cockroach NavPas channel resolved in the closed state revealed an interaction between the CTD EF-hand and the alpha helix within the III-IV inactivation loop (Shen et al., 2017). A prominent feature of the CTD:loop contact is a hydrophobic interaction between Phe-1501 on the EF-hand with Tyr-1136 and Ala-1139 on the inactivation loop helix (Figure 3a). The EF-hand on the CTD of Na_v_1.5 displays 53% amino acid sequence conservation to that of NavPas, including aliphatic Na_v_1.5 Leu-1861 in place of the NavPas Phe-1501. Structural alignment-based substitution of the Na_v_1.5 EF-hand into the CTD:loop complex shows a likely hydrophobic interaction of Leu-1861 with Ala-1139, which is conserved in the Na_v_1.5 loop helix as Ala-1496 (Figure 3b). To test whether L1861 plays a significant role in Na_v_1.5 gating, we analyzed the properties of Na_v_1.5^L1861A^ compared to Na_v_1.5^S1786ter^ and wild-type channels. Voltage-dependent steady-state inactivation of Na_v_1.5^L1861A^ had V_1/2_ = −114.6 +/- 1.5 mV, an 11.5 mV hyperpolarizing shift from wild-type Na_v_1.5 (V_1/2_ = −103.1 +/- 0.6 mV), approaching the voltage shift of Na_v_1.5^S1786ter^ to V_1/2_ = −116.8 +/- 1.3 mV (Figure 2a, Table 1). The L1861A substitution also strongly accelerated the rate of closed-state inactivation to a degree approaching that of Na_v_1.5^S1786ter^ (Figure 2b, Table 1). However, unlike Na_v_1.5^S1786ter^, the Na_v_1.5^L1861A^ mutation did not discernably alter the rate of open-state inactivation or current density, and had only a small effect on persistent current (Figure 2c, Table 1). While Na_v_1.5^L1861A^ also showed a right-shift in the apparent voltage of channel activation to −37 mV (Figure 2e), the voltage dependence of activation for inactivation-defective Na_v_1.5[ID]^L1861A^ did not significantly differ from Na_v_1.5[ID] (Figure 2f, Table 1), thereby showing that the L1861A mutation does not directly alter the activation gating mechanism.

**Figure 3.**
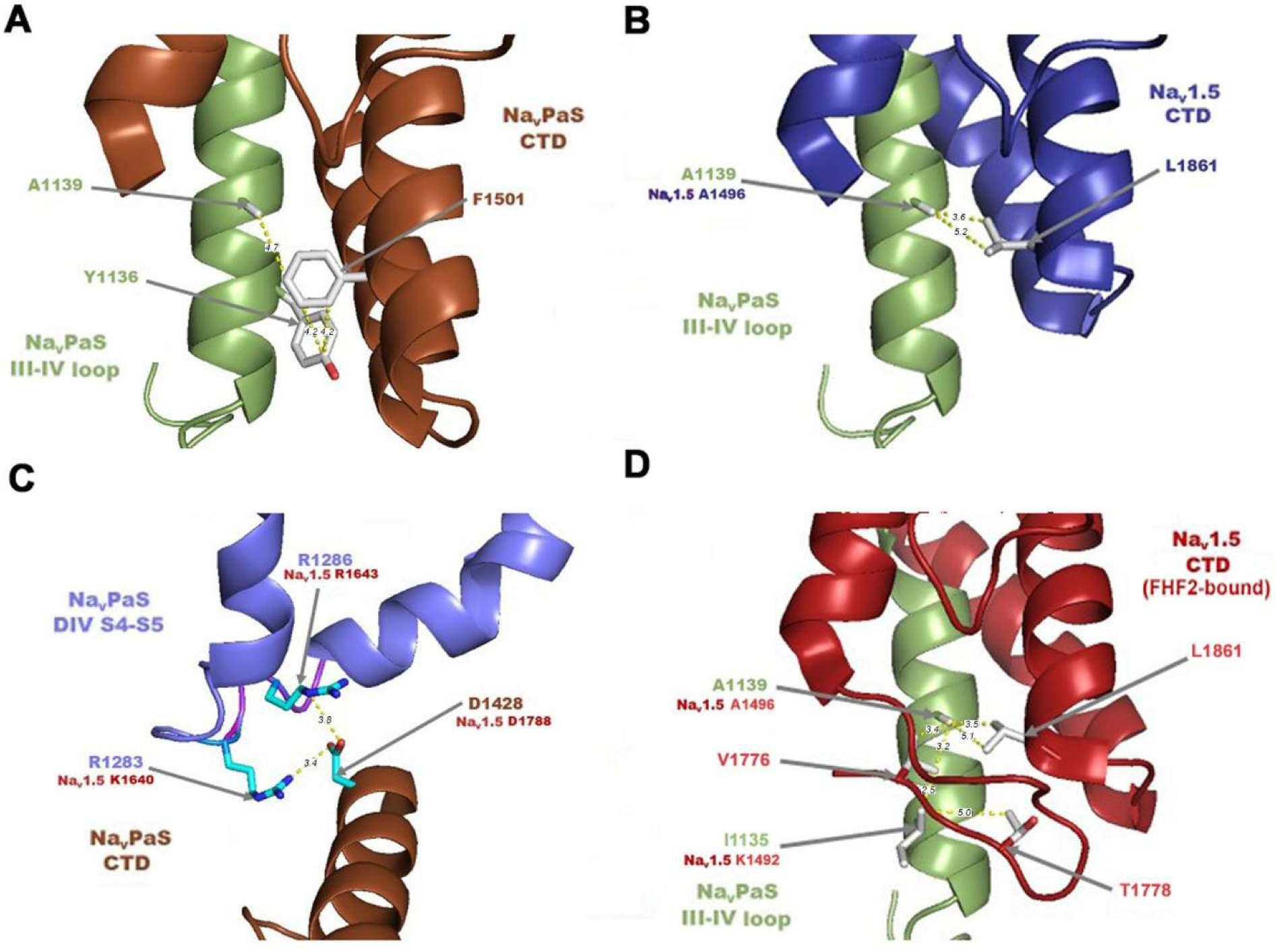
Structural models of intramolecular sodium channel CTD interactions guiding selection of missense mutations for electrophysiological analysis. **(A)** Interaction of the NavPas CTD with the III-IV inactivation loop showing a hydrophobic contact (with Angstrom distances indicated) between the CTD residue F1501 and the III-IV inactivation loop residues A1139 and Y1136. (from structure 5X0M). **(B)** Hypothetical interaction of the Na_v_1.5 CTD with the III-IV inactivation loop generated by substituting the Na_v_1.5 CTD (from structure 4OVN) in place of the NavPas CTD from panel A, and showing the hypothetical hydrophobic interaction of CTD L1861 with NavPas loop A1139 (A1496 in Na_v_1.5) in the III-IV loop. **(C)** Interaction of the NavPas CTD with the DIV S4/S5 helix showing an electrostatic interaction between CTD residue D1428 (D1788 in Na_v_1.5) and the DIV S4/S5 helix residues R1283 and R1286 (K1640 and R1643 in Na_v_1.5). **(D)** Hypothetical enhanced interaction of the Na_v_1.5 CTD with the III-IV inactivation loop upon FHF2B binding to the CTD (from structure 4DCK). The β-strand conformation adopted by the JM region brings the Na_v_1.5 CTD residues V1776 and T1778 in close proximity to the Na_v_Pas III-IV loop residues I1135 (K1492 in Na_v_1.5) and A1139 (A1496 in Na_v_1.5).

Many FHF proteins induce depolarizing shifts to the voltage dependence of Nav steady-state inactivation (Goldfarb, 2024). Consistent with previous findings (Wang et al., 2011b), FHF2B induced an 11 mV depolarizing shift in V_1/2_ inactivation (V_1/2_ = −92.0 +/- 1.7 mV) (Figure 4a, Table 2). Since FHF2B can still bind to Na_v_1.5 bearing the L1861A mutation (Suppl. Fig.1), we tested whether FHF2B can still modulate Na_v_1.5^L1861A^ inactivation. Indeed, FHF2B depolarized the V_1/2_ inactivation of Na_v_1.5^L1861A^ by 9 mV (V_1/2_ = −105.8 +/- 1.8 mV)(Figure 4a, Table 2). Therefore, while the L1861A mutation alters intrinsic inactivation gating, the mutation does not significantly interfere with the ability of FHF2B to modulate channel inactivation.

**Figure 4.**
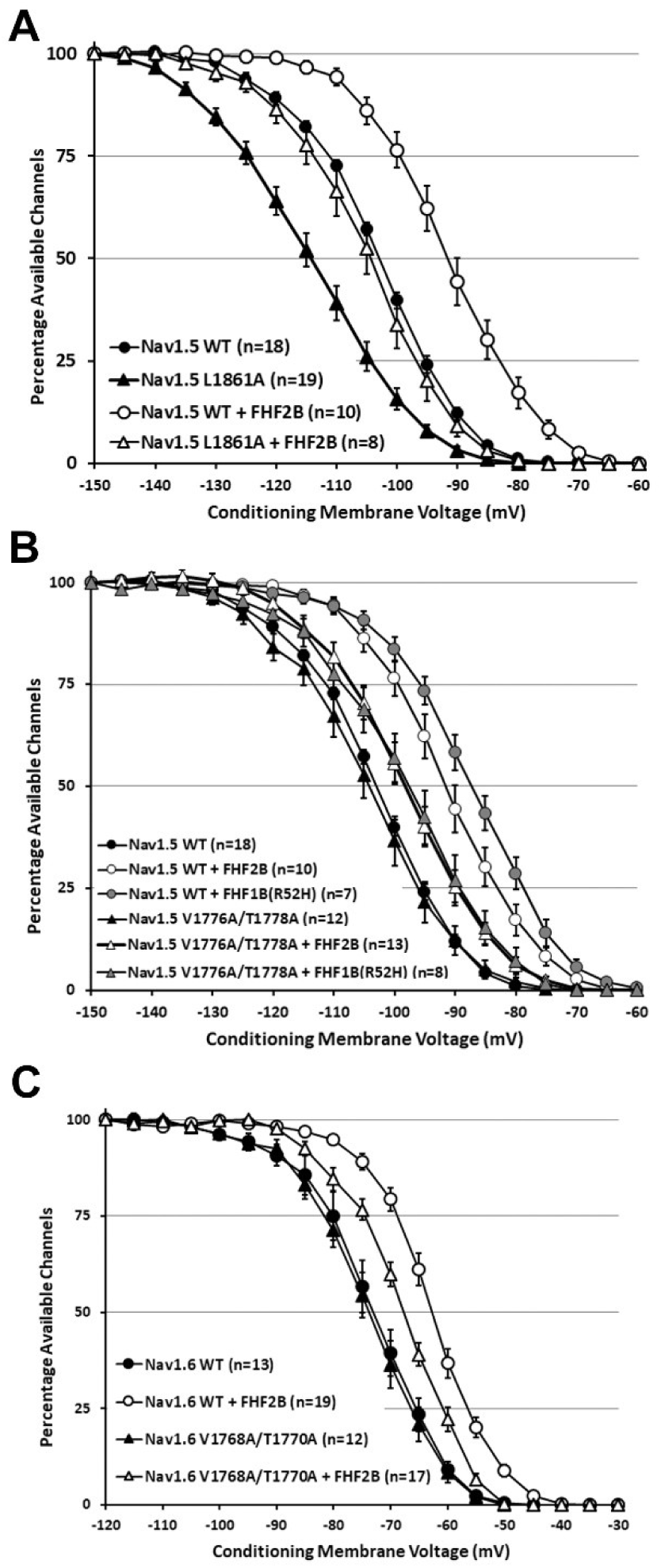
FHF modulation of wild-type and mutant sodium channels. **(A)** FHF2B induces an 11 mV depolarizing shift in steady-state inactivation of Na_v_1.5^wildtype^ and a 9 mV depolarizing shift for inactivation of Na_v_1.5^L1861A^. **(B)** In the absence of FHF, voltage dependence of Na_v_1.5^V1776A/T1778A^ inactivation is indistinguishable from that of Na_v_1.5^wildtype^. However, depolarizing modulation of channel inactivation by either FHF2B or FHF1B^R52H^ is significantly reduced for Na_v_1.5^V1776A/T1778A^ when compared to Na_v_1.5^wildtype^. **(C)** In the absence of FHF, the voltage dependence of Na_v_1.6^V1768A/T1770A^ is indistinguishable from that of Na_v_1.6^wildtype^. However, depolarizing modulation of channel inactivation by FHF2B is significantly reduced for Na_v_1.6^V1768A/T1770A^ when compared to Na_v_1.6^wildtype^. For all assays, the number of cells (n) measured for each transfection is indicated within the figure panels.

**Table 2.** FHF effects on gating of Na_v_1.5 and Na_v_1.6 wild-type and mutant derivatives. Parameter values for mutant channels that significantly differ from those of corresponding wild-type channels along with parameter values for channels significantly modulated by FHFs are shown in bold italics. ^a^P<0.0001 for effect of FHF on a channel’s gating. ^b^P<0.02 for effect of FHF on channel’s gating and P<0.007 for reduction of FHF gating effect on the JM mutant vs. wild-type channel.

| hNav1.5 +/- FHF Voltage Dependence of Steady-State Inactivation |  |  |  |  |  |  |  |  |
| --- | --- | --- | --- | --- | --- | --- | --- | --- |
|  | Wild-Type |  |  | V1776A/T1778A |  |  | L1861A |  |
|  | w/o FHF | + FHF2B | + FHF1B<sup>R52H</sup> | w/o FHF | + FHF2B | + FHF1B<sup>R52H</sup> | w/o FHF | + FHF2B |
| V<sub>1/2</sub> (mV) | -103.1 +/- 0.6 | **-92.0 +/- 1.7<sup>a</sup>** | **-87.0 +/- 1.9<sup>a</sup>** | -104.4 +/- 1.8 | **-98.4 +/- 1.4<sup>b</sup>** | **-97.9 +/- 2.1<sup>b</sup>** | -114.6 +/- 1.5 | **-105.8 +/- 1.8<sup>a</sup>** |
| k (mV) | 7.2 +/- 0.2 | 6.7 +/- 0.2 | 7.2 +/- 0.4 | 7.8 +/- 0.3 | 7.3 +/- 0.2 | 7.7 +/- 0.4 | 9.1 +/- 0.2 | **6.8 +/- 0.2<sup>a</sup>** |
| # Cells | 18 | 10 | 7 | 12 | 13 | 8 | 19 | 8 |

| hNav1.5 +/- FHF Voltage Dependence of Activation |  |  |  |  |  |  |  |  |
| --- | --- | --- | --- | --- | --- | --- | --- | --- |
|  | Wild-Type |  |  | V1776A/T1778A |  |  | L1861A |  |
|  | w/o FHF | + FHF2B | + FHF1B<sup>R52H</sup> | w/o FHF | + FHF2B | + FHF1B<sup>R52H</sup> | w/o FHF | + FHF2B |
| V<sub>1/2</sub> (mV) | -52.9 +/- 0.9 | -50.7 +/- 0.7 | -48.5 +/- 0.8 | -50.4 +/- 1.7 | -54.9 +/- 2.0 | -51.4 +/- 1.6 | **-37.1 +/- 1.5<sup>a</sup>** | -47.7 +/- 1.5 |
| k (mV) | 5.4 +/- 0.2 | 4.6 +/- 0.3 | 4.3 +/- 0.3 | 5.9 +/- 0.4 | 5.5 +/- 0.5 | 5.1 +/- 0.3 | **10.5 +/- 0.3<sup>a</sup>** | 5.7 +/- 0.7 |
| # Cells | 16 | 8 | 6 | 7 | 5 | 5 | 13 | 5 |

| mNav1.6<sup>TTXr</sup> +/- FHF2B Voltage Dependence of Steady-State Inactivation |  |  |  |  |
| --- | --- | --- | --- | --- |
|  | Wild-Type |  | V1768A/T1770A |  |
|  | w/o FHF | + FHF2B | w/o FHF | + FHF2B |
| V<sub>1/2</sub> (mV) | -72.8 +/- 1.7 | **-62.2 +/- 0.8<sup>a</sup>** | -73.7 +/- 1.5 | **-67.7 +/- 1.0<sup>b</sup>** |
| k (mV) | 6.4 +/- 0.1 | 5.4 +/- 0.2 | 6.5 +/- 0.4 | 6.0 +/- 0.4 |
| # Cells | 13 | 20 | 12 | 17 |

TABLE 2. FHF effects on gating of Nav1.5 and Nav1.6 wild-type and mutant derivatives
| mNav1.6<sup>TTXr</sup> +/- FHF Voltage Dependence of Activation |  |  |  |  |
| --- | --- | --- | --- | --- |
|  | Wild-Type |  | V1768A/T1770A |  |
|  | w/o FHF | + FHF2B | w/o FHF | + FHF2B |
| V<sub>1/2</sub> (mV) | -20.0 +/- 0.9 | -23.4 +/- 0.9 | -17.5 +/- 1.0 | -18.1 +/- 1.0 |
| k (mV) | 6.9 +/- 0.2 | 5.4 +/- 0.2 | 7.7 +/- 0.3 | 6.8 +/- 0.1 |
| # Cells | 10 | 9 | 9 | 6 |

### EF-hand Missense Mutant Na_v_1.5^D1788K^ Partially Phenocopies the Activation Gating and Persistent Current Abnormalites of Na_v_1.5^S1786ter^

The structure of the cockroach NavPas channel resolved in the closed state also revealed an electrostatic interaction between the sidechains of Asp-1428 in the CTD EF-hand and Arg-1283 and Arg-1286 in Domain IV on the cytoplasmic face of the channel (Figure 3c)(Shen et al., 2017). These residues are conserved in Na_v_1.5 (Asp-1788 in CTD, Lys-1640 and Arg-1643 in Domain IV), suggesting a similar CTD interaction. Na_v_1.5^D1788K^ was constructed and its gating properties were analyzed. The transient current amplitude and the voltage dependence of inactivation for Na_v_1.5^D1788K^ were statistically indistinguishable from those of wild-type Na_v_1.5 (Figure 2a, Table1). However, Na_v_1.5^D1788K^ generated greatly elevated persistent current (Figure 2d, Table 1).

Na_v_1.5^D1788K^ also displayed a strong depolarizing shift in the voltage dependence of channel activation compared to wild-type Na_v_1.5 (V_1/2_ = −38.1 +/- 1.3 mV) (Figure 2e, Table 1). As the D1788K substitution did not significantly alter the inactivation properties of Na_v_1.5, it seemed plausible that this substitution directly affects the activation gating mechanism. As confirmation, an inactivation-defective derivative of Na_v_1.5^D1788K^ was constructed and analyzed. Indeed, Na_v_1.5[ID]^D1788K^ exhibited a significant depolarizing shift in the voltage dependence of activation (V_1/2_ = 45.0 +/- 0.9mV) compared to Na_v_1.5[ID] (V_1/2_ = 50.5 +/- 1.1 mV) (Figure 2f, Table 1).

### The JM Segment Missense Mutant Na_v_1.5^V1776A/T1778A^ and Paralogous Mutant Na_v_1.6^V1768A/T1770A^ Impair FHF Modulation of Inactivation Gating

Crystal structural analysis of the Na_v_1.5 CTD has shown that FHF2B binding does not alter the conformation of EF-hand and α6 segments of the CTD, and causes only an 8° bend of the hinge between these segments (Wang et al., 2012; Wang et al., 2014). However, the JM region undergoes dramatic restructuring in the presence of FHF2B. Closer examination of this FHF-associated structure showed that Na_v_1.5 JM residues SerValAlaThreGlu (1775-1779) adopt a β-strand conformation that interfaces with the EF-hand in such a way as to increase the potential interaction surface with the III-IV inactivation loop helix (Figure 3d). The hypothetical alignment of the FHF2B-bound Na_v_1.5 CTD with the inactivation loop helix reveals multiple potential hydrophobic interactions of JM Val-1776 and Thr-1778 with the inactivation loop (Figure 3d). If CTD:loop interaction dictates the voltage setpoint for channel inactivation by restraining the transition from closed to inactivated state, then FHF restructuring of the JM segment that enhances loop interaction could serve as a basis for FHF modulation of inactivation. Guided by these crystal structures, we engineered Na_v_1.5^V1776A/T1778A^ and analyzed its inactivation properties in the absence and presence of FHF.

The voltage dependence of Na_v_1.5^V1776A/T1778A^ steady-state inactivation was indistinguishable from that of wild-type Na_v_1.5 in the absence of FHF (Figure 4b, Tables 1,2)(V_1/2_ −104.4 +/- 1.8 mV vs. - 103.1 +/- 0.6 mV, *P* > 0.8). However, FHF2B modulation of Na_v_1.5^V1776A/T1778A^ inactivation was significantly impaired. While FHF2B induced an 11 mV depolarizing shift in the V_1/2_ of steady-state inactivation for wild-type Na_v_1.5, FHF2B only shifted the V_1/2_ of inactivation for Na_v_1.5^V1776A/T1778A^ by 6 mV (V_1/2_ = −98.4 +/- 1.4 mV)(Figure 4b, Table 2). FHF2B still interacts with the mutant channel, as shown by coimmunoprecipitation (Suppl Fig 1). Na_v_1.5^V1776A/T1778A^ also showed impaired modulation by FHF1B^R52H^, which shifts V_1/2_ of inactivation for wild-type Na_v_1.5 by 17 mV to −87.2 +/- 1.7 mV but only shifts V_1/2_ of Na_v_1.5^V1776A/T1778A^ by 6 mV to −98.4 +/- 1.6 mV (Figure 4b, Table 2). Therefore, the Na_v_1.5 JM segment acts as a generalized effector of inactivation gating modulation by FHFs.

The JM segment sequences of vertebrate sodium channels are highly conserved, with the valine and threonine residues described above found in all VGSCs except Na_v_1.9. Therefore, we tested whether these two residues contribute to FHF modulation of another VGSC, the neuronal Na_v_1.6. Wild-type Na_v_1.6 or Na_v_1.6^V1768A/T1770A^ were transiently expressed in Neuro2A cells in the absence or presence of FHF2B. FHF2B induced a 10 mV depolarizing shift in steady-state inactivation for wild-type Na_v_1.6, from V_1/2_ = −72.8 +/- 1.7 mV without to V_1/2_ = −62.6 +/- 0.8 mV in the presence of FHF2B (Figure 4c, Table 2). In contrast, FHF2B only induced a 6 mV depolarizing shift in for Na_v_1.6^V1768A/T1770A^, from V_1/2_ = −73.7 +/- 1.5 mV without to V_1/2_ = −67.7 +/- 1.0 mV (Figure 4c, Table 2). These findings suggest a generalized role for the juxtamembrane segment in contributing to FHF modulation of VGSC inactivation gating.

## Discussion

We have shown that the cytoplasmic carboxyl-terminal domain (CTD) of cardiac Na_v_1.5 modulates many properties of the channel, including: 1) voltage-dependent closed-state inactivation, 2) rate of inactivation from the closed state, 3) rate of inactivation from the open state, 4) FHF binding, 5) FHF-dependent modulation of closed-state inactivation, 6) voltage-dependent activation, 7) persistent sodium current amplitude, and 8) transient sodium current amplitude. The EF-hand fold within the CTD influences all of these properties, as shown by the contrasting phenotypes of Na_v_1.5^S1786ter^ versus Na_v_1.5^E1900ter^. Within the EF-hand fold, Leu-1861 is critical for establishing the proper setpoint and rate of closed-state inactivation, while making little or no contribution to other CTD properties, surprisingly including open-state inactivation rate. Another EF-hand residue, Asp-1788, is required for properly setting the voltage for channel activation and for helping suppress persistent current, while exerting little or no effect on inactivation gating or magnitude of transient sodium current. Within the Na_v_1.5 JM segment, Val-1776 and/or Thr-1778 serve as an effector for FHF modulation of voltage-dependent inactivation without observable effects on intrinsic properties of the channel.

Cryo-EM structures of full-length sodium channels (Shen et al., 2017; Yan et al., 2017b) along with crystal structures of the Na_v_1.5 CTD +/- FHF2 (Wang et al., 2012; Wang et al., 2014) guided our selection of mutant constructs and support several potential mechanisms underlying mutational effects on channel gating. The interactions of the III-IV inactivation loop helix with the CTD in the closed state or with the cytoplasmic base of the pore in the inactivated state are mutually incompatible due to steric constraints, thereby demanding dissociation of the CTD from the inactivation loop during inactivation (Shen et al., 2017; Yan et al., 2017b). Voltage sensor movements associated with membrane depolarization must either drive the dissociation of the CTD from the loop or expose a higher affinity site at the base of the pore for the loop to dock. Mutations predicted to disrupt or diminish CTD interaction with the III-IV inactivation loop helix would thereby facilitate closed-state inactivation at more negative potentials, as shown here for Na_v_1.5^L1861A^, and as shown previously for Na_v_1.5^E1783K^ (Gade et al., 2020). Building upon this hypothesis, we reasoned that FHF-induced depolarizing shifts to the voltage dependence of channel inactivation may result from an expansion of the contact surface of the CTD with the III-IV inactivation loop helix. Crystal structures obtained for the Na_v_1.5 in the absence vs. presence of complexed FHF2B revealed an FHF-induced conformational change in the Na_v_1.5 JM segment (Wang et al., 2012; Wang et al., 2014) in a manner that could increase the contact surface of the CTD with the III-IV loop helix (Figure 2). Consistent with this mechanism of FHF functionality, mutation of highly conserved JM segment residues in Na_v_1.5 or Na_v_1.6 specifically impaired FHF-induced depolarizing voltage shifts for channel inactivation without altering intrinsic inactivation gating of the channels.

The closed-state structure of NavPas (Shen et al., 2017) also revealed an electrostatic interaction between an acidic residue in the CTD EF-hand and two basic residues in the IVS4/S5 helical region at the cytoplasmic base of the channel, with all three of these residues conserved in mammalian sodium channels (Figure 2c). The corresponding EF-hand mutated cardiac channel Na_v_1.5^D1788K^ displayed a marked depolarizing shift in the voltage dependence of activation without obvious effects on channel inactivation. The basic residue sidechains of Lys-1640 and Arg-1643 predicted to interact with Na_v_1.5 Asp-1788 (Shen et al., 2017) are at the carboxyl-terminal end of the Na_v_1.5 domain IV voltage sensor (R1622/R1625/R1628/R1631/R1634/R1637/*<u>K1640</u>*/*<u>R1643</u>*) (Yan et al., 2017b). The shielding of these sensor charges by the CTD may modestly reduce the inward force exerted by membrane potential on the voltage sensor, allowing wild-type Na_v_1.5 to open at more negative potential than Na_v_1.5^D1788K^.

The structural basis by which persistent sodium current is minimized following inactivation remains poorly understood due to a lack of CTD resolution in reported structures of inactivated VGSCs. Prior work demonstrated that either calmodulin or FHF must be bound to the CTD in order to suppress persistent current (Abrams et al., 2020). We have shown here that Na_v_1.5^D1788K^ displays elevated persistent current. This finding raises the possibility that the CTD remains electrostatically latched to the domain IV S4/S5 region after inactivation mediated by the domain III-IV loop. By doing so, the CTD in conjunction with associated FHF or calmodulin may create a steric barrier restricting inactivation loop dissociation and minimizing persistent current.

## Acknowledgments

We thank Hanna Cloutier for excellent assistance with electrophysiology recordings and Flopateer Shenouda for assistance with plasmid constructions. This work was supported by NIH grant R01 HL142498 and by City University of New York Research Foundation award 7F303.

---

**Supplemental Figure 1.**
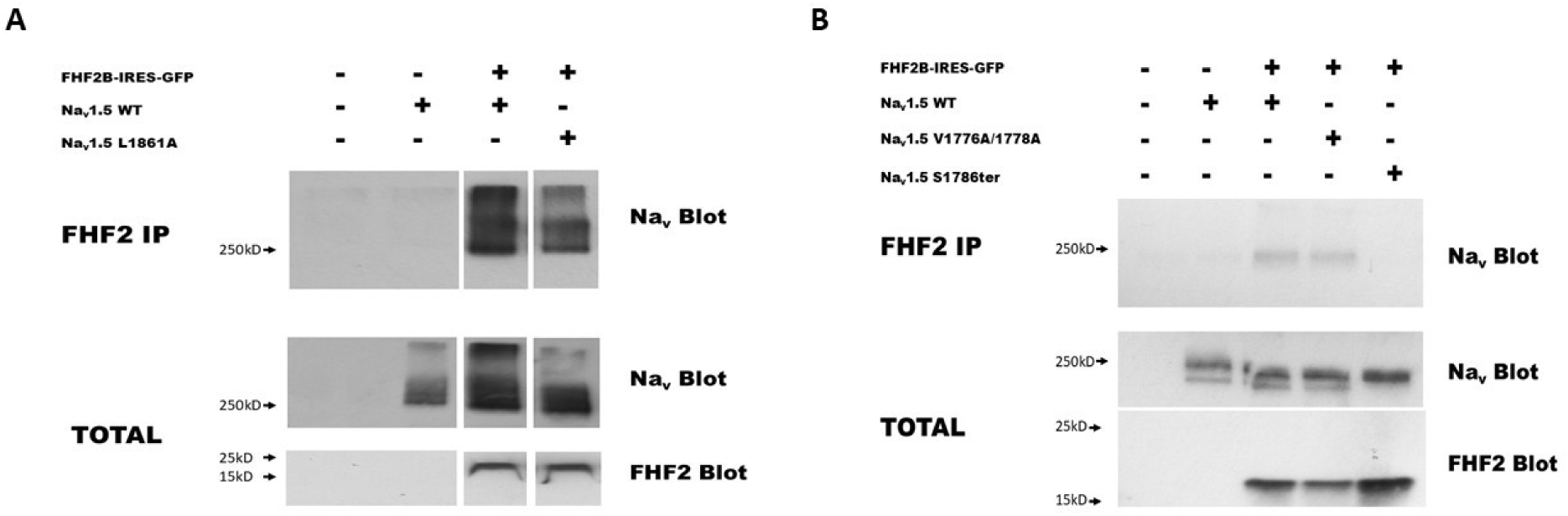
Na_v_1.5 CTD mutations L1861A and V1776A/T1778A do not disrupt FHF2B binding. **(A)** Na_v_1.5^L1861A^ binds FHF2B in co-transfected cells. N2a cells were transiently transfected with expression vectors as indicated at top. Total lysates or anti-FHF2 immunoprecipitates (IP) were subjected to gel electrophoresis and immunoblotting with anti-pan-Nav or anti-FHF2. Na_v_1.5 is detected in anti-FHF2 IPs from cells expressing FHF2B and either Na_v_1.5 WT (wild-type) or Na_v_1.5^L1861A^. Cells without FHF2B were cotransfected with empty vector pIRES-GFP. **(B)** Na_v_1.5^V1776A/T1778A^ binds FHF2B in co-transfected cells. Na_v_1.5 is detected in anti-FHF2 IPs from cells expressing FHF2B and either Na_v_1.5 WT (wild-type) or Na_v_1.5^V1776A/T1778A^, but not in cells expressing FHF2B and CTD deletion mutant Na_v_1.5^S1786ter^.

## Notes

### Competing Interest Statement

The authors have declared no competing interest.

